# Re-introduction of endogenous pathways for propionyl-CoA, 1-propanol and propionate formation in *Escherichia coli*

**DOI:** 10.1101/2021.11.21.469472

**Authors:** Thai Le

## Abstract

In a previous study, we found that 2-ketobutyrate (2-KB) was seriously degraded in *Escherichia coli*. In the present investigation, we tried to clarify the products of that degradation process, and intriguingly reconfirmed that 2-KB is chopped up to form propionyl-CoA, 1-propanol and propionate. This short commentary re-introduces efficient endogenous pathways for production of value-added odd-chain compounds such as propionyl-CoA-derived chemicals.

## 1. Summary of propionyl-CoA, 1-propanol and propionate formation in *Escherichia coli* in literature

Acetyl-CoA is a key intermediate for even-chain compound production, while propionyl-CoA plays that role in case of odd-chain products. Unfortunately, propionyl-CoA is a non-native metabolite to most organisms (Srirangan et al., 2017), and usually therefore, propionate is exogenously added for propionyl-CoA formation. From that propionyl-CoA, several value-added compounds, such as 1-propanol and propionate, have been produced.

To supply propionyl-CoA for synthesis of a high-valuable polyketide, namely 6-deoxyerythronolide B (6-dEB) in *E. coli*, propionate was exogenously added and endogenous propionyl-CoA ligase (PrpE) was over-expressed (Pfeifer et al., 2001) (Fig. 1A). Later, the same group adopted an endogenous sleeping beauty mutase (Sbm) pathway (discussed below) for propionyl-CoA formation, but this pathway subsequently was found to be inefficient and expensive, requiring cyanocobalamin (vitamin B_12_) (Zhang et al., 2010). In another study, formation of an odd-chain acid, 3-hydroxyvalerate, was attempted via propionyl-CoA, and again propionyl-CoA ligase (PrpE from *Salmonella typhimurium* LT2) or propionyl-CoA transferase (Pct from *Megasphaera elsdenii*) was utilized with propionate addition (Martin et al., 2013) (Fig. 1B).

**Figure 1.**
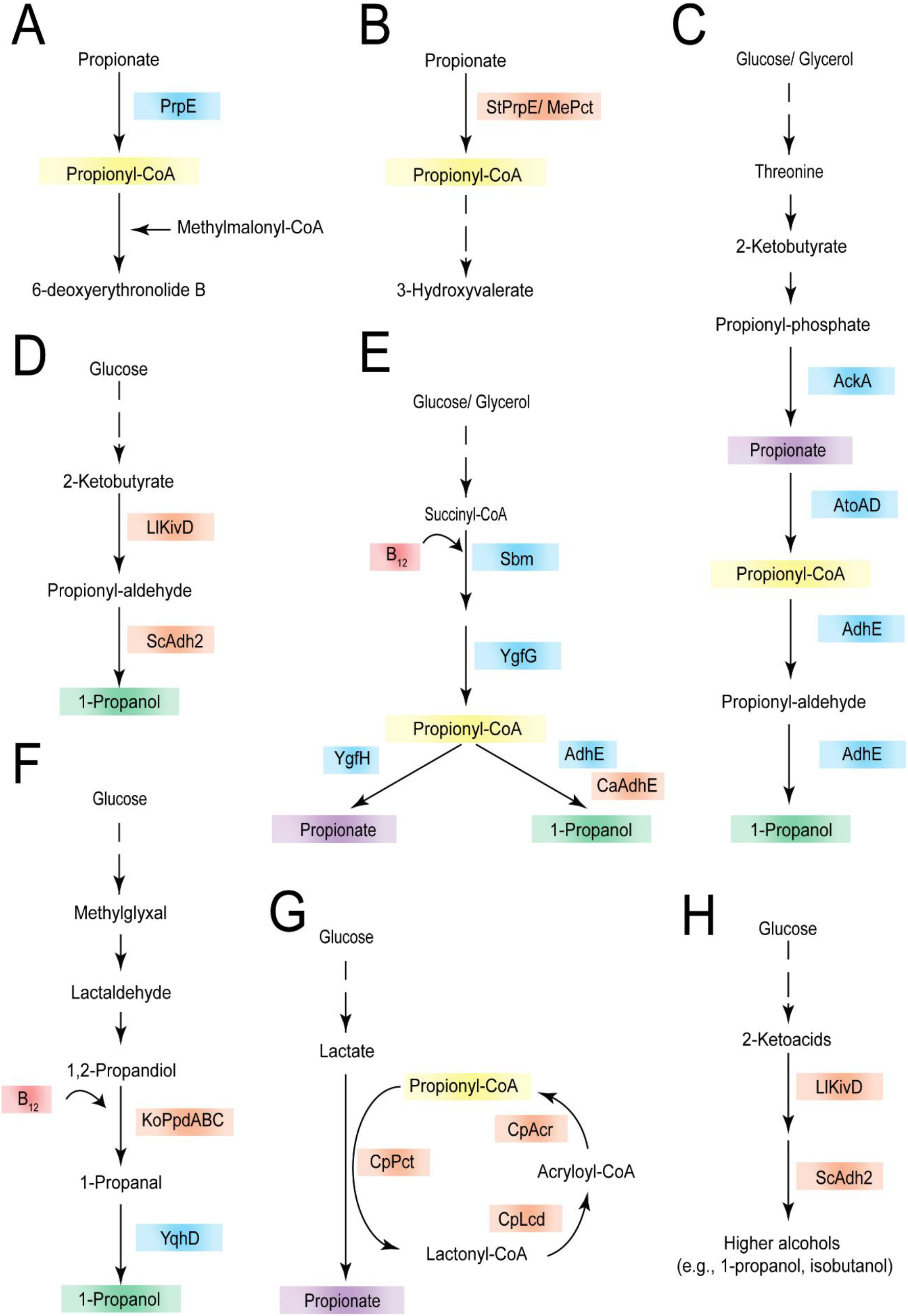
Summary of propionyl-CoA, 1-propanol and propionate formation in *Escherichia coli* in literature. **(A)** Propionyl-CoA formation from propionate by endogenous PrpE for polyketide, 6-deoxyerythronolide B (6-dEB) synthesis (Pfeifer et al., 2001). **(B)** Propionyl-CoA formation from propionate by exogenous StPrpE or MePct for 3-hydroxyvalerate production (Martin et al., 2013). **(C)** 1-Propanol production via 2-KB by AckA, AtoAD and AdhE (Jun Choi et al., 2012). **(D)** Production of 1-propanol via 2-KB by LlKivd and ScAdh2 (Shen and Liao, 2008; 2013). **(E)** Utilization of endogenous sleeping beauty mutase (Sbm) pathway for 1-propanol and propionate production (Srirangan et al., 2014). **(F)** 1-Propanol production via 1,2-propanediol by KoPpdABC and YqhD (Jain and Yan, 2012). **(G)** Propionate production by employment of acrylate pathway genes from *Clostridium propionicum* (Kandasamy et al., 2013). **(H)** Higher-alcohol production via 2-ketocaids (Atsumi et al., 2008). **Abbreviations: PrpE**, propionyl-CoA ligase; **Pct**, propionyl-CoA transferase; **AckA**, acetate kinase A; **AtoAD**, acetyl-CoA:acetoacetyl-CoA synthase; **AdhE**, alcohol dehydrogenase; **Kivd**, 2-ketoacid decarboxylase; **Adh2**, alcohol dehydrogenase 2; **Sbm**, sleeping beauty mutase/ methylmalonyl-CoA mutase; **YgfG**, methylmalonyl-CoA decarboxylase; **YgfH**, propionyl-CoA:succinate transferase; **PpdABC**, diol dehydratase; **YqhD**, alcohol dehydrogenase; **Lcd**, lactoyl-CoA dehydratase; **Acr**, acryloyl-CoA reductase; **St**, *Salmonella typhimurium;* **Me**, *Megasphaera elsdenii*; **Ll**, *Lactococcus lactis*; **Sc**, *Saccharomyces cerevisiae*; **Ca**, *Clostridium acetobutylicum;* **Ko**, *Klebsiella oxytoca;* **Cp**, *Clostridium propionicum* **Solid line**, one-step reaction; **Dashed line**, multi-step reaction. Yellow boxes, propionyl-CoA; Green boxes, 1-propanol; Purple boxes, propionate; Blue boxes, homologous enzymes; Orange boxes, heterologous enzymes

Regarding 1-propanol, Choi *et al*. engineered *E. coli* for 1-propanol production from glucose and glycerol via 2-ketobutyrate (2-KB) via a complex pathway. In that research, 2-KB was subsequently converted to propionyl-phosphate, propionate, and propionyl-CoA with overexpression of AckA and AtoAD. Eventually, 1-propanol was formed from propionyl-CoA by AdhE via the propionyl-aldehyde intermediate (Jun Choi et al., 2012) (Fig. 1C). In other research, a heterologous alcohol dehydrogenase 2 (Adh2 from *Saccharomyces cerevisiae*) and a promiscuous 2-ketoacid decarboxylase (Kivd from *Lactococcus lactis*) were introduced into *E. coli* to obtain 1-propanol (Shen and Liao, 2008; 2013a) (Fig. 1D). By attempting the endogenous Sbm pathway, Srirangan et al. could concomitantly produce propionate and 1-propanol in *E. coli*. However, this pathway was determined to be inherently weak and expensive, since an exogenous supply of cyanocobalamin (vitamin B_12_) was needed (Srirangan et al., 2013). After >200h cultivation, only ~7g/L 1-propanol and >1g/L propionate were recorded, but large amounts of other products, specifically ethanol (>30g/L), lactate, acetate (9g/L), and succinate (~2g/L) were formed (Srirangan et al., 2014) (Fig. 1E). In another endeavor, the same group produced >10g/L propionate, but the already-noted limitations, which is to say, byproduct formation and expense (supply of vitamin B_12_), remained (Akawi et al., 2015). For 1-propanol production, Jian *et al*., by another approach, expanded a 1,2-propanediol route via introduction of a heterologous diol dehydratase into *E. coli*. Nevertheless, the pathway was unfavorable (three heterologous genes were needed, and toxic intermediate methylglyoxal accumulated) as well as expensive (exogenous vitamin B_12_ had to be added) (Jain & Yan, 2011; Jain et al., 2015) (Fig. 1F).

In an effort to produce propionate in *E. coli*, Kandasamy *et al*. employed the acrylate pathway from *Clostridium propionicum*, but the performane was poor, the titer as low as 4 mM, and large amounts of other products appeared (Kandasamy et al., 2013) (Fig. 1G). And once again (see above), the Sbm pathway was adopted for propionate formation (Srirangan et al., 2014) (Fig. 1E).

2-Keto acids have been confirmed to be promising intermediates for production of higher alcohols (Atsumi et al., 2008) (Fig. 1H). By adopting the promiscuous 2-keto acid decarboxylase (Kivd) and heterologous alcohol dehydrogenase 2 (Adh2), several alcohols, from straight-chain alcohols to branched-chain and aromatic counterparts, were produced. However, among those, only isobutanol achieved industrial-scale titer and yield (~22g/l and 86% theoretical yield; Atsumi et al., 2008) due mainly to the high activity of Kivd with 2-ketoisovalerate (De La Plaza et al., 2004). Later, the same group attempted to produce 1-propanol and/or butanol by utilizing the above strategy, but the titer and yield were low (Shen and Liao, 2008; 2013a) (Fig. 1D). These results indicate the importance of a good enzyme for decarboxylation of 2-keto acids to produce non-native higher alcohols. We recommend, as alternatives to the same Kivd for all 2-keto acids, pyruvate dehydrogenase (PDH) for 1-propanol production and branched-chain 2-keto acid dehydrogenases for 1-butanol, 2-methyl-1-butanol and 3-methyl-1-butanol formation (Atsumi et al., 2008) (Fig. 1H).

## 2. Propionyl-CoA, 1-propanol and propionate formation via 2-KB degradation

In our previous study, AceE, PoxB and PflB were identified as being responsible for degradation of 2-KB (Le and Park, 2021). The degradation of 2-KB, therefore, was hypothesized to yield propionyl-CoA (then 1-propanol) and propionate. By comparing the chromatogram of 2-KB degradation after 24h with those of the 1-propanol and propionate standards (Fig. 2), we successfully confirmed that hypothesis. The synthesis of propionyl-CoA, propionate and 1-propanol in *E. coli* W3110 by endogenous enzymes, therefore, is re-introduced in Fig. 3 below. The 2-KB is synthesized from threonine by IlvA (aerobically) or TdcB (anaerobically). Then, it is oxidized to propionate by PoxB, or converted to propionyl-CoA by PDH (under the aerobic condition) or PflB (under the anaerobic condition). Propionyl-CoA can be further converted, again, to propionate by Pta-AckA and/or 1-propanol by AdhE. Additionally, 2-KB can form D-2-hydroxybutyrate (D-2-HB) by LdhA, but this enzyme (reaction) seemingly is less active. In contrast, the conversion of D/L-2-HB to 2-KB is favorable (Le and Park, 2021).

**Figure 2.**
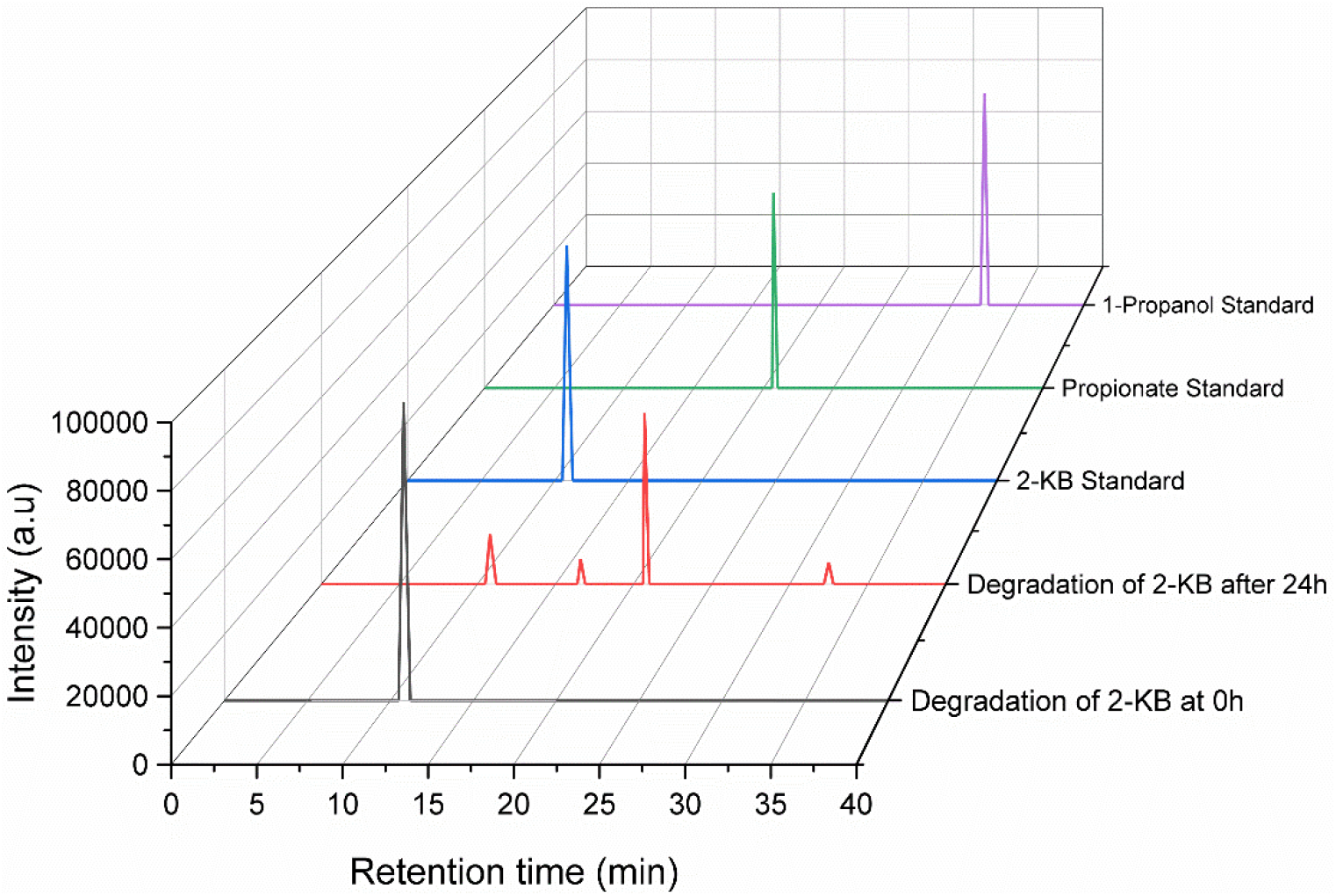
Confirmation of degradation of 2-KB to propionate and 1-propanol in wild-type *E. coli* W3110 by high-performance liquid chromatography (HPLC)

**Figure 3.**
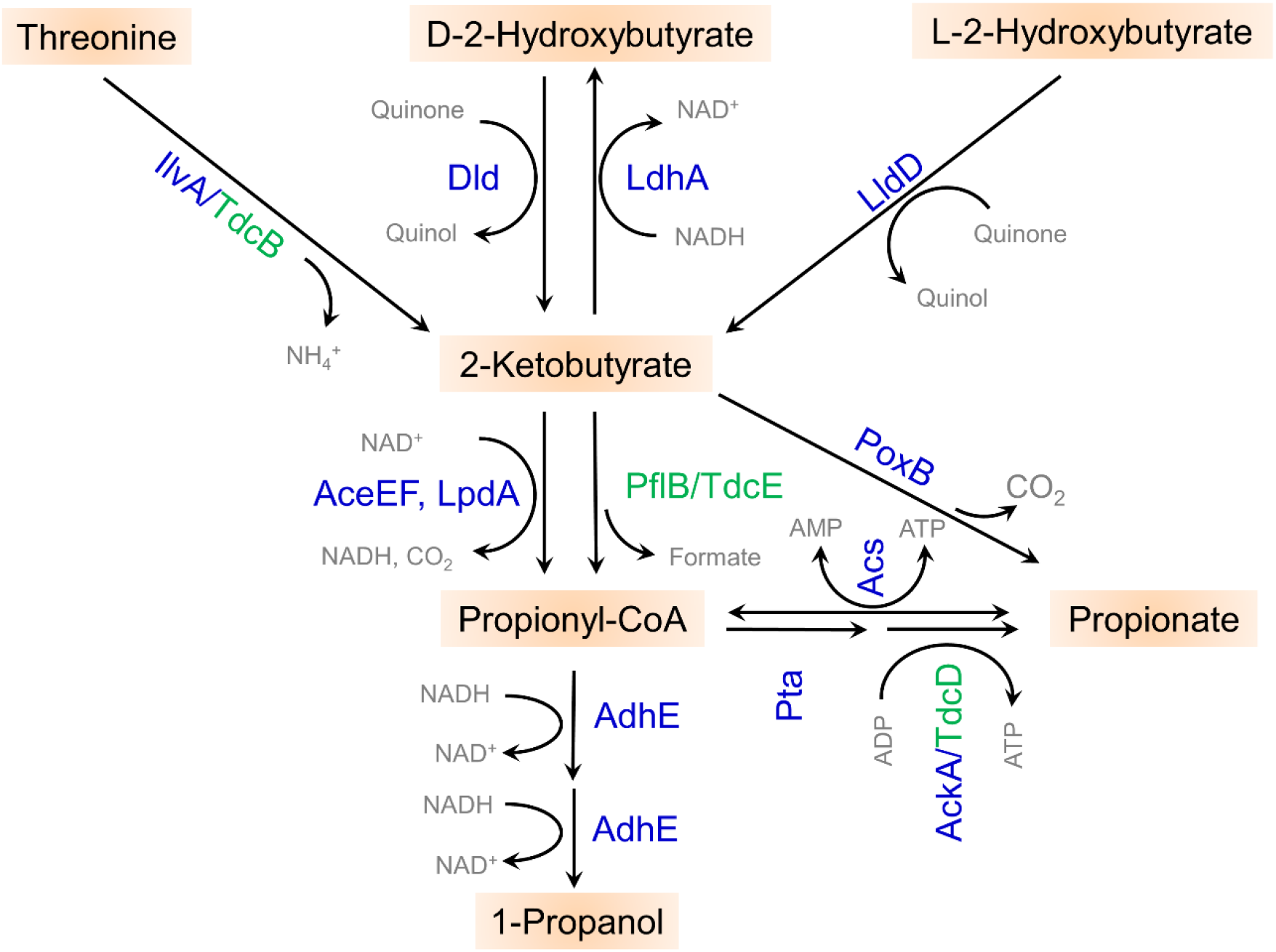
Metabolism of 2-KB to propionyl-CoA, 1-propanol and propionate in *E. coli*. **Abbreviations: IlvA,** Threonine dehydratase; **TdcB**, Threonine dehydratase; **Dld**, quinone-dependent D-Lactate dehydrogenase; **LdhA**, NADH-dependent D-Lactate dehydrogenase; **LldD**, quinone-dependent L-Lactate dehydrogenase; **AceE**; Pyruvate dehydrogenase (PDH) subunit 1; **AceF**, PDH subunit 2; **LpdA**, PDH subunit 3; **PflB**, Pyruvate-formate lyase; **TdcE** Ketoacid-formate lyase; **PoxB**, Pyruvate oxidase; **Pta**, Phosphate acetyltransferase; **AckA**, Acetate kinase; **TdcD**; Propionate kinase; **Acs**, Acetyl-coenzyme A synthetase; **AdhE**, Aldehyde-alcohol dehydrogenase. Enzymes in blue are proposed to function aerobically, and those in green, anaerobically.

To our best knowledge, Bisswanger was the first to report that PDH exhibits very good activity towards 2-KB *in vitro* (Bisswanger, 1981), which was later adopted for production of 3-hydroxyvalerate from a propionate-unrelated carbon source (Tseng et al., 2010) and odd-chain fuels/chemicals (Tseng and Prather, 2012). Nonetheless, the good activity of PDH to 2-KB seemingly has not been well recognized, due to the paucity of studies, as discussed above, that have utilized it for propionyl-CoA intermediate formation and/or propionate and 1-propanol production.

Intriguingly, 1-propanol formation from threonine was reported recently based on an investigation of the growth of *E. coli* as surface-attached biofilms, and the metabolic pathway for that formation was proposed as well. However, there might have been some misunderstanding. In that study, propionyl-CoA was proposed to derive from 2-KB via PflB and TdcE under the aerobic and anaerobic conditions respectively, but we believe that the enzymes should have been actually PDH (AceEF, LpdA) and PflB/TdcE. Besides, our present review helps to further elucidate the degradation of threonine in *E. coli* (Heßlinger et al., 1998).

## 3. Over-expression of AdhE for 1-propanol production from 2-KB

Propionate and 1-propanol formation from 2-KB was examined by means of a resting cell of *E. coli* W3110. AdhE for 1-propanol formation (NADH-consuming reaction) (Fig. 3) and *Cb*FDH (formate dehydrogenase [FDH] from *Candida boidinii* (Le and Park, 2021)) for NADH regeneration were over-expressed, and investigations were conducted both aerobically and anaerobically.

Under the aerobic condition (i.e., +O_2_), both recombinants (AdhE and AdhE+FDH) performed similarly. Namely, most of the 2-KB was converted to propionate, which was foreseen (Fig. 4). This was due to the fact that FDH was intentionally over-expressed for NADH regeneration to accelerate the 1-propanol formation reaction, while the NADH was strongly competed against by the electron transport chain, as witnessed previously (Le and Park, 2021). As a result, no appreciable difference between those two recombinants was seen.

**Figure 4.**
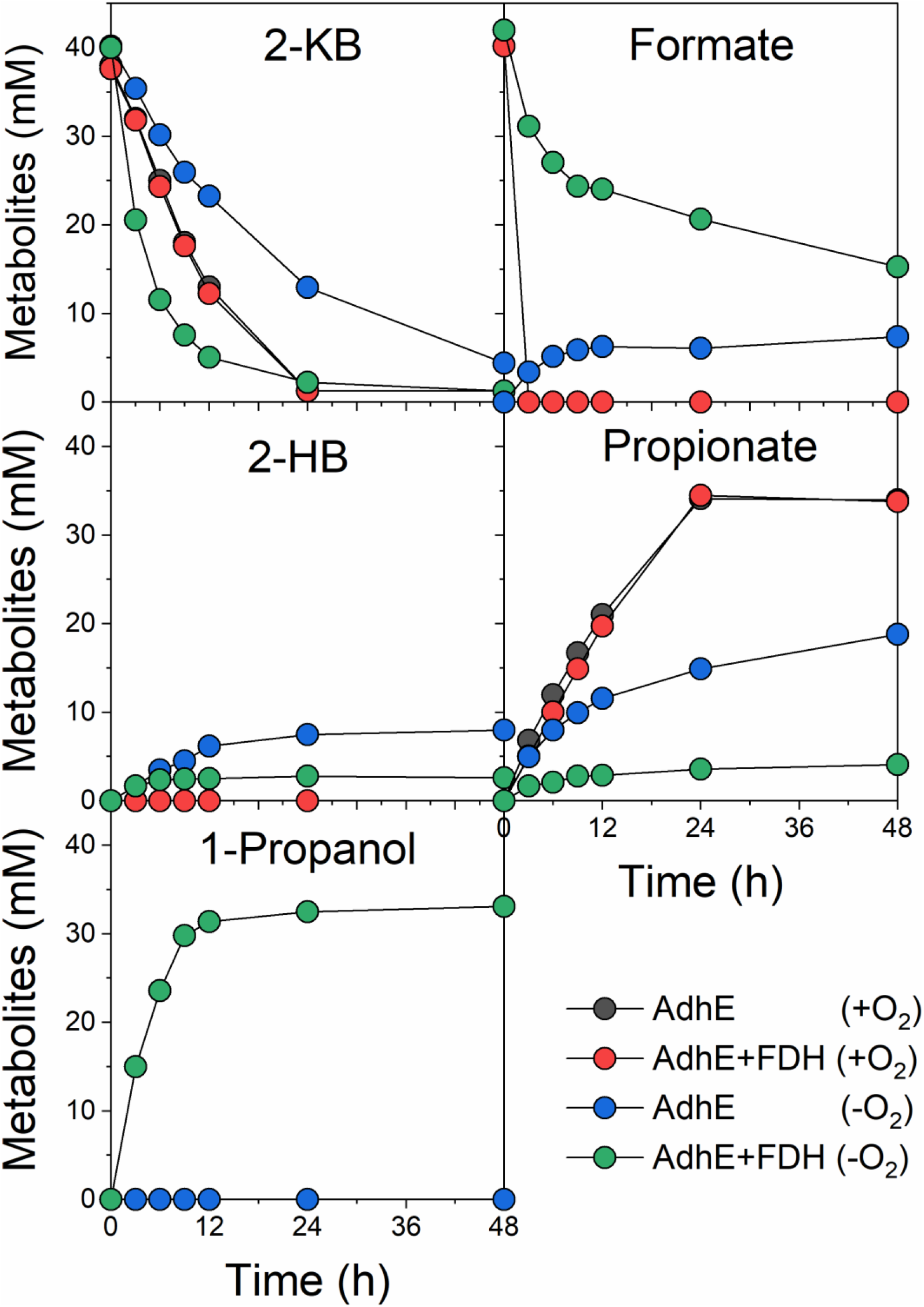
Over-expression of AdhE for 1-propanol production from 2-KB by resting cell of *E. coli* W3110. **AdhE**, *E. coli* W3110 with over-expression of AdhE; **AdhE+FDH**, *E. coli* W3110 with over-expression of AdhE and *Cb*FDH; **+O_2_**, study was conducted aerobically; **−O_2_**, study was conducted anaerobically

Under the anaerobic condition (i.e., −O_2_), the expression of AdhE alone could not accelerate 1-propanol production, because two NADHs would be needed for this reaction (Fig. 3). With the help of *Cb*FDH for generation of NADH, most of the 2-KB was diverted to 1-propanol, as expected (AdhE+FDH (−O_2_)).

This study again confirmed the endogenous pathways for propionyl-CoA, 1-propanol and propionate formation from 2-KB, which are depicted in Fig. 3 above.

## 4. Conclusion

The formation of propionyl-CoA, 1-propanol, and propionate from 2-KB by homologous enzyme in *E. coli* was re-examined. The benefits of 2-KB metabolism for high-value propionyl-CoA-derived chemical production was reconfirmed. This utility is owed to the fact that for even-chain compound production, the metabolism and regulation of a vital intermediate, acetyl-CoA, have been studied intensively while those of propionyl-CoA, for odd-chain products, lag behind. This paper re-introduces efficient endogenous routes that hopefully can accelerate value-added propionyl-CoA-derived compound production. By adopting those routes, neither exogenous toxic substrate (i.e., propionate)/expensive cofactor (i.e., vitamin B_12_) nor low-efficient heterologous enzymes are needed.

## 5. Materials and methods

### 5.1. Materials

The hosts and materials (with sources in parentheses) were the same as utilized in Le and Park, 2021: *Escherichia coli* DH5 alpha (Takara (Korea)); primers and sequence confirmations (Macrogen Co. Ltd (Korea)); chromosomal isolation kit (Promega (USA)); DNA purification kits (Qiagen (Germany)); DNA polymerase (Invitrogen (Korea)); restriction enzymes and T4 ligase (New England Biolabs (USA)); yeast extract, agar and tryptone (Difco (USA)); all of the other chemicals were procured from Sigma-Aldrich (USA), unless indicated otherwise.

### 5.2. Methods

#### 5.2.1. Plasmid construction for overexpression of *Ec_AdhE*

To construct pB (harboring *adhE*) plasmid, *Ec_adhE* gene was PCR-amplified from the chromosomal DNA of *E. coli* W3110. Then, the PCR product was cloned into pQE-80L (Qiagen (Germany)). The pA (harboring *Cb_fdh*) was the same as utilized in Le and Park, 2021.

#### 5.2.2. Bioconversion of 2-KB to 1-propanol

Non-growing-cell bioconversion was conducted in 100 mM potassium phosphate buffer (PPB (pH 7.0)) at an agitation speed of 200 rpm and a temperature of 30°C, while the density was 5.0 OD_600_ (otherwise indicated), and 2-KB and formate were ~40 mM. To generate the anaerobic condition, helium gas was purged into the headspace of the bottle (closed by rubber cap).

#### 5.2.3. Analytical methods

Please refer to Le and Park, 2021. Briefly, formate, 2-KB, 2-HB, propionate and 1-propanol were detected by HPLC with 2.5 mM H_2_SO_4_ as the mobile phase.

## Abbreviation

2-KB: 2-ketobutyrate
2-HB: 2-hydroxybutyrate
FDH: formate dehydrogenase

## References

Akawi, L., Srirangan, K., Liu, X., Moo-Young, M., Perry Chou, C., 2015. Engineering Escherichia coli for high-level production of propionate. J. Ind. Microbiol. Biotechnol. 42, 1057–1072. https://doi.org/10.1007/s10295-015-1627-4

Atsumi, S., Hanai, T., Liao, J.C., 2008. Non-fermentative pathways for synthesis of branched-chain higher alcohols as biofuels. Nature 451, 86–89. https://doi.org/10.1038/nature06450

Bisswanger, H., 1981. Substrate specificity of the pyruvate dehydrogenase complex from Escherichia coli. J. Biol. Chem. 256, 815–822. https://doi.org/10.1016/s0021-9258(19)70050-7

De La Plaza, M., Fernández De Palencia, P., Peláez, C., Requena, T., 2004. Biochemical and molecular characterization of α-ketoisovalerate decarboxylase, an enzyme involved in the formation of aldehydes from amino acids by Lactococcus lactis. FEMS Microbiol. Lett. 238, 367–374. https://doi.org/10.1016/j.femsle.2004.07.057

Heßlinger, C., Fairhurst, S.A., Sawers, G., 1998. Novel keto acid formate-lyase and propionate kinase enzymes are components of an anaerobic pathway in Escherichia coli that degrades L-threonine to propionate. Mol. Microbiol. 27, 477–492. https://doi.org/10.1046/j.1365-2958.1998.00696.x

Jain, R., Sun, X., Yuan, Q., Yan, Y., 2015. Systematically Engineering Escherichia coli for Enhanced Production of 1,2-Propanediol and 1-Propanol. ACS Synth. Biol. 4, 746–756. https://doi.org/10.1021/sb500345t

Jain, R., Yan, Y., 2011. Dehydratase mediated 1-propanol production in metabolically engineered Escherichia coli. Microb. Cell Fact. 10, 97. https://doi.org/10.1186/1475-2859-10-97

Jun Choi, Y., Hwan Park, J., Yong Kim, T., Yup Lee, S., 2012. Metabolic engineering of Escherichia coli for the production of 1-propanol. Metab. Eng. 14, 477–486. https://doi.org/10.1016/j.ymben.2012.07.006

Kandasamy, V., Vaidyanathan, H., Djurdjevic, I., Jayamani, E., Ramachandran, K.B., Buckel, W., Jayaraman, G., Ramalingam, S., 2013. Engineering Escherichia coli with acrylate pathway genes for propionic acid synthesis and its impact on mixed-acid fermentation. Appl. Microbiol. Biotechnol. 97, 1191–1200. https://doi.org/10.1007/s00253-012-4274-y

Le, T., Park, S., 2021. Development of efficient microbial cell factory for whole-cell bioconversion of L-threonine to 2-hydroxybutyric acid. Bioresour. Technol. 126090. https://doi.org/10.1016/j.biortech.2021.126090

Martin, C.H., Dhamankar, H., Tseng, H.C., Sheppard, M.J., Reisch, C.R., Prather, K.L.J., 2013. A platform pathway for production of 3-hydroxyacids provides a biosynthetic route to 3-hydroxy-γ-butyrolactone. Nat. Commun. 4, 1–10. https://doi.org/10.1038/ncomms2418

Pfeifer, B.A., Admiraal, S.J., Gramajo, H., Cane, D.E., Khosla, C., 2001. Biosynthesis of Complex Polyketides in a Metabolically Engineered Strain of E. coli. Science (80-.). 291, 1790–1793.

Shen, C.R., Liao, J.C., 2013. Synergy as design principle for metabolic engineering of 1-propanol production in Escherichia coli. Metab. Eng. 17, 12–22. https://doi.org/10.1016/j.ymben.2013.01.008

Shen, C.R., Liao, J.C., 2008. Metabolic engineering of Escherichia coli for 1-butanol and 1-propanol production via the keto-acid pathways. Metab. Eng. 10, 312–320. https://doi.org/10.1016/j.ymben.2008.08.001

Srirangan, K., Akawi, L., Liu, X., Westbrook, A., Blondeel, E.J.M., Aucoin, M.G., Moo-Young, M., Chou, C.P., 2013. Manipulating the sleeping beauty mutase operon for the production of 1-propanol in engineered Escherichia coli. Biotechnol. Biofuels 6, 1–14. https://doi.org/10.1186/1754-6834-6-139

Srirangan, K., Bruder, M., Akawi, L., Miscevic, D., Kilpatrick, S., Moo-Young, M., Chou, C.P., 2017. Recent advances in engineering propionyl-CoA metabolism for microbial production of value-added chemicals and biofuels. Crit. Rev. Biotechnol. 37, 701–722. https://doi.org/10.1080/07388551.2016.1216391

Srirangan, K., Liu, X., Westbrook, A., Akawi, L., Pyne, M.E., Moo-Young, M., Chou, C.P., 2014. Biochemical, genetic, and metabolic engineering strategies to enhance coproduction of 1-propanol and ethanol in engineered Escherichia coli. Appl. Microbiol. Biotechnol. 98, 9499–9515. https://doi.org/10.1007/s00253-014-6093-9

Tseng, H.C., Harwell, C.L., Martin, C.H., Prather, K.L.J., 2010. Biosynthesis of chiral 3-hydroxyvalerate from single propionate-unrelated carbon sources in metabolically engineered E. coli. Microb. Cell Fact. 9, 1–12. https://doi.org/10.1186/1475-2859-9-96

Tseng, H.C., Prather, K.L.J., 2012. Controlled biosynthesis of odd-chain fuels and chemicals via engineered modular metabolic pathways. Proc. Natl. Acad. Sci. U. S. A. 109, 17925–17930. https://doi.org/10.1073/pnas.1209002109

Zhang, H., Boghigian, B.A., Pfeifer, B.A., 2010. Investigating the role of native propionyl-CoA and methylmalonyl-CoA metabolism on heterologous polyketide production in Escherichia coli. Biotechnol. Bioeng. 105, 567–573. https://doi.org/10.1002/bit.22560

